# Bridging-mediated compaction of mitotic chromosomes

**DOI:** 10.1101/2022.09.27.509716

**Authors:** Giada Forte, Lora Boteva, Nick Gilbert, Peter R. Cook, Davide Marenduzzo

**Affiliations:** SUPA, School of Physics and Astronomy, University of Edinburgh, Peter Guthrie Tait Road, Edinburgh, EH9 3FD, UK; MRC Human Genetics Unit, Institute of Genetics and Cancer, University of Edinburgh, Western General Hospital, Crewe Road South, Edinburgh, EH4 2XU, UK; Sir William Dunn School of Pathology, University of Oxford, Oxford, OX1 3RE, England, UK

## Abstract

Eukaryotic chromosomes compact during mitosis and meiosis into elongated cylinders – and not the spherical globules expected of self-attracting long flexible polymers. This process is mainly driven by condensin-like proteins. Here, we present Brownian-dynamics simulations involving two types of such proteins. The first anchors topologically-stable and long-lived chromatin loops to create bottlebrush structures. The second forms multivalent bridges between distant parts of these loops without entrapping them. We show bridging factors lead to the formation of shorter and stiffer mitotic-like cylinders, without requiring any energy input. These cylinders have several features matching experimental observations. For instance, the axial condensin backbone breaks up into clusters as found by microscopy, and cylinder elasticity qualitatively matches that seen in chromosome pulling experiments. Additionally, simulating global condensin depletion or local faulty condensin loading gives phenotypes in agreement with experiments, and provides a mechanistic model to understand mitotic chromatin structure at common fragile sites.

## INTRODUCTION

During mitosis and meiosis, chromosomes condense to form the iconic cylinders seen by light microscopy (1; 2). Understanding how such cylinders form is a fundamental question which has not yet been fully answered. Compaction into a cylinder instead of a sphere is surprising from the perspective of polymer physics, as polymers subjected to self-attraction usually collapse into spherical globules (3). Experiments show that fibre condensation is mediated by SMC (structural maintenance of chromosome) proteins like condensins and cohesins, and disentanglement by topoisomerases (4). While the contour length of loops remains unchanged as cells pass through mitosis (5), rearranging them to form a long string of consecutive chromatin loops creates a bottlebrush polymer with a large effective stiffness or persistence length that is a prerequisite of cylindrical mitotic structures (2; 6). Surprisingly, histone proteins, which are essential constituents of chromatin, are not required for cylinder formation (7). As both condensins and topoisomerases are ATP-dependent (4), it is also normally assumed that active processes are required for condensation.

The condensins and SMC proteins that play such central roles in mitosis organise chromosomes locally in two distinct ways: by topologically loading onto fibres to stabilise long-lived loops (8; 9), and by bridging two different genomic segments (without embracing either) to drive clustering with other bound SMC proteins (10; 11). There are also two types of condensin that play different roles in mitosis: condensin II binds during prophase to form an axial scaffold, while condensin I is cytoplasmic and binds during metaphase to shorten chromosomes (9; 12; 13; 14). At the global level, the topologically associating domains (TADs), seen in interphase using a high-throughput variant of chromosome-conformation capture (3C) known as Hi-C, are typically lost during mitosis in minutes, as the condensin II axial backbone winds up into a helix (9). Notwithstanding this, imaging shows the gross structure of interphase chromosome territories is preserved into metaphase (15).

The loop-extrusion model, assuming that condensins are motors moving on chromatin and extruding genomic loops, provides an appealing way to explain how the ATP-dependent activity of condensins can be harnessed to both generate and stabilise loops anchored to the backbone of a mitotic chromosome (6). Steric exclusion between different loops attached to the backbone then creates a large persistence length, which can be far greater than that of the underlying chromatin fibre, and is a possible reason for the cylindrical appearance of chromosomes during mitosis and meiosis (2; 16; 17). While extremely useful as a starting point, this model still leaves many open questions. For example, the role of condensin-mediated bridging (rather than looping) is not directly addressed, and it remains unclear how further bottlebrush compaction might occur and whether it requires energy. Super-resolution and micromanipulation experiments suggest that condensins self-assemble into relatively inhomogeneous columns inside mitotic chromosomes (18), and the reason for this is unclear. Additionally, the elasticity of human metaphase chromosomes is striking as they can be stretched tenfold by an external force, and yet they relax back to their original length once the force is removed (19).

Here, we develop and characterise a simple polymer model to study chromosome compaction. Significantly, our simulations do not involve constraining the polymer in a cylinder, as often done previously; then, the resulting shape emerges solely from specified interactions. Our work includes two types of condensin-like proteins: one stabilises loops (which provide an underlying bottlebrush geometry), and another binds multivalently to chromatin to form local bridges and clusters without requiring energy input. We suggest such bridging becomes particularly relevant after prophase, when cytoplasmic condensin I associates with chromatin. We do not wish to suggest that condensin I solely acts as a bridge, but given its enhanced concentration at the onset of metaphase, we argue that bridging by the newly added condensin I drives the striking morphological transition from the prophase bottlebrush to a shorter and stiffer mitotic cylinder. This compaction depends on the statistics and size of loops and topoisomerase activity. We also simulate the response of these structures to stretching, finding that the qualitative behaviour is similar to that observed experimentally. Finally, we provide new insights into folding around common fragile sites (CFSs) – genomic regions of up to ∼1.2 Mbp in which chromosomal lesions often appear following replication stress (20). Our simple model complements simulations of chromosome folding based on loop extrusion (6; 9), and points to the crucial importance of condensin-mediated bridging in chromosome self-assembly.

## RESULTS

### A polymer model for mitotic chromosome folding

We study the formation of mitotic chromosomes via coarse-grained molecular dynamics (MD) simulations (Fig. 1). A schematic of the model used is shown in Fig. 1. As experiments suggest mitotic chromosomes are arranged in consecutive loops (8; 21; 22; 23) with contour lengths much the same as found during interphase (5), we begin with a looped polymer depicted as a relaxed bottlebrush (Fig. 1, left). Loops in this polymer do not change during simulations; we imagine their molecular anchors are provided by SMC proteins, and that this loop configuration is created by active (6) or diffusive (24) loop extrusion. However, our focus is on the later folding dynamics driven by condensin bridging [Fig. 1, right and (9; 13; 12; 14; 18; 25)].

**Fig. 1:**
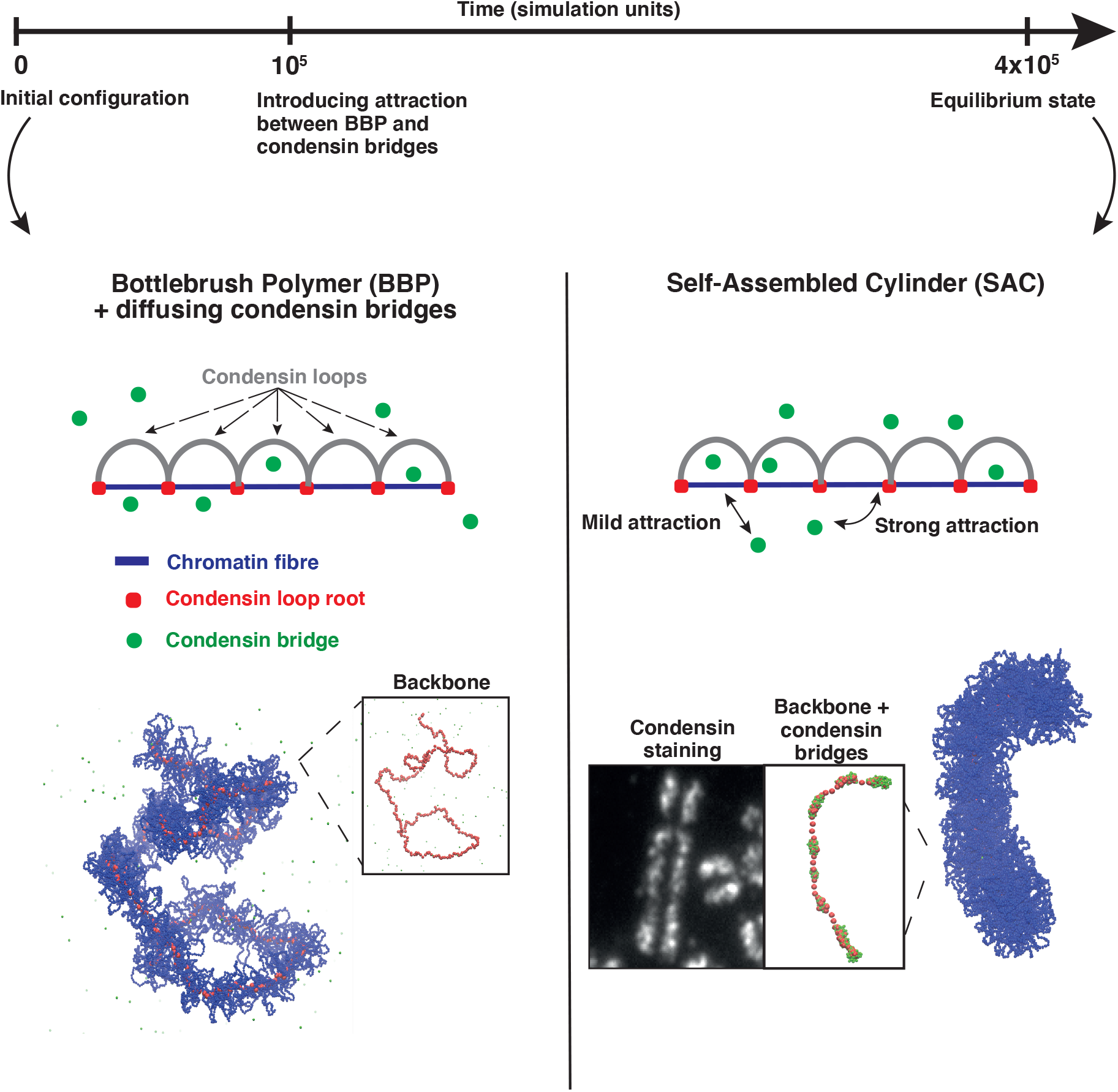
Simulations of mitotic chromosomes. (Top) Simulation timeline. (Centre) Sketch of the model ingredients. Left: a mitotic prophase chromosome (blue line) is modelled as a bottlebrush polymer (BBP), with consecutive condensin loops (gray arcs, modelled as springs). Condensin loop anchors and condensin bridges are shown as red and green dots respectively; the latter experience a purely steric interaction with the polymer in the first part of the simulation. Right: starting from 10^5^*τ*_*LJ*_ condensin bridges can bind reversibly to chromatin, weakly to blue beads, and strongly to red ones, generating a self-assembled cylinder (SAC). (Bottom) Snapshots from computer simulations showing typical structures for the bottlebrush polymer (left) and self-assembled cylinder (right) regimes. The transition between the two is driven by condensin-mediated bridging. From left to right the three insets correspond to: backbone (condensin loop anchors) in the bottlebrush regime, experimental condensin staining in metaphase, and backbone with condensin bridges in the self-assembled cylinder regime. The experimental condensin staining reveals an inhomogenous profile as emerges from our simulations.

We take into consideration cases where either all condensin-mediated loops have the same contour length *L*_*loop*_, or contour length varies following a Poisson distribution with average value *L*_*loop*_. We consider *L*_*loop*_ = 40, 50, 60, *σ*, where the bead diameter *σ* corresponds to the fibre diameter which is typically 10-30nm, and each bead is assumed to contain 2kbp (see Star Methods). These loop sizes are realistic, as mitotic loops are usually 80-120 kbp long (8; 5; 26). Condensin bridges are modelled as diffusing beads (shown in green in Fig. 1) that bind reversibly and weakly to chromatin blue beads and strongly to red beads – the loop anchors (see Star Methods for force field used). Typically, numbers of condensin bridges and loops are comparable, although exact numbers do not qualitatively affect results.

The underlying chromatin fibre is characterized by a persistence length *l*_*p*_ = 3*σ* ∼ 60 nm which is consistent with that of interphase chromatin (27). To model topoisomerase activity simply, chromatin strand passing is allowed as pairs of non-bonded polymer beads interact via a soft potential

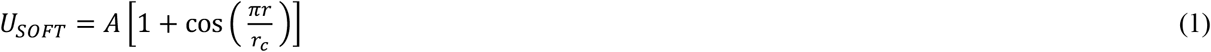

where *r* is the distance between two beads and *r*_*c*_ = 2^1/6^*σ* is a cut-off distance such that *U*_*SOFT*_ = 0 when *r*> *r*_*c*_. The thermal energy of the system is *k*_*B*_*T* where *k*_*B*_ is the Boltzmann constant and *T* is the temperature. Therefore, when *A* = 1 *k*_*B*_*T*, polymer beads can cross each other because of thermal fluctuations, effectively modelling catalysis by topoisomerase II of strand cutting, movement of another strand through the cut, and ligation. Increasing *A* to 100 *k*_*B*_*T* models a reduction in topoisomerase activity, as in other works (8; 28).

### Condensin-mediated bridging compacts bottlebrushes into cylinders

We first consider the case where all loops have the same contour length in an initial (pre-equilibrated) prophase-like state – the bottlebrush-polymer (BBP) configuration in Fig. 1, left. Starting from the BBP, attractive interactions are switched on between condensin bridges and chromatin. A striking morphological transition now occurs (Suppl. Movie 1): the bottlebrush (Fig. 1, left) tightens up and becomes significantly shorter and stiffer, forming a structure reminiscent of a metaphase chromosome (Fig. 1, right). We refer to this final configuration as a self-assembled cylinder (SAC). Compaction into a cylinder is driven by bridging, as bridges (like condensins) are assumed to be multivalent and able to bind more than one chromatin bead (or loop anchor). More specifically, the bridging-induced attraction (29) provides a general mechanism to cluster bridges and to compact the polymer. It is based on positive feedback: bridging increases binding-site concentration locally, which recruits further bridges, and this triggers clustering. This would collapse a simple unlooped polymer into a spherical globule (29), but here competition between the bridging-induced compaction and looping-induced stiffening of the BPP drives self-assembly into cylinders.

Condensins in SACs are concentrated in local clusters scattered along axial columns– as seen in mitotic chromosomes *in vivo* [(18), and Fig. 1, right inset]. We suggest the non-uniform axial distribution is formed as a central condensin column breaks up in an effect akin to the Rayleigh instability: bridging-induced clustering creates an effective surface tension, so when the interfacial energy becomes too large to maintain a contiguous column/stream, the column/stream breaks up into smaller globules. Loop size, *L*_*loop*_, and soft repulsion, *A*, affect cluster size: the larger either is, the smaller clusters are (Fig. S1).

We next quantify the geometric changes as the BBP morphs into a SAC in two ways (Fig. 2A). First, the average gyration radius, *R*_*g*_, was measured (Figs. 2A and 2Bi, top). *R*_*g*_ sharply decreases once condensin binding begins (at *t* = 0 in Fig. 2Bi, top); this is in accord with biological observations (2). Interestingly, the extent of compaction depends on *L*_*loop*_ and *A*: increasing *L*_*loop*_ increases *R*_*g*_, while *R*_*g*_ increases with decreasing topoisomerase II activity (Fig. 2Bi, top). Consequently, strong topoisomerase activity (when *A* becomes comparable to thermal energy) leads to more compaction. These results are consistent with the intuition that longer loops and stronger repulsion yield larger excluded volumes preventing compaction, and exemplify another important role of topoisomerase II in chromosome folding.

**Fig. 2:**
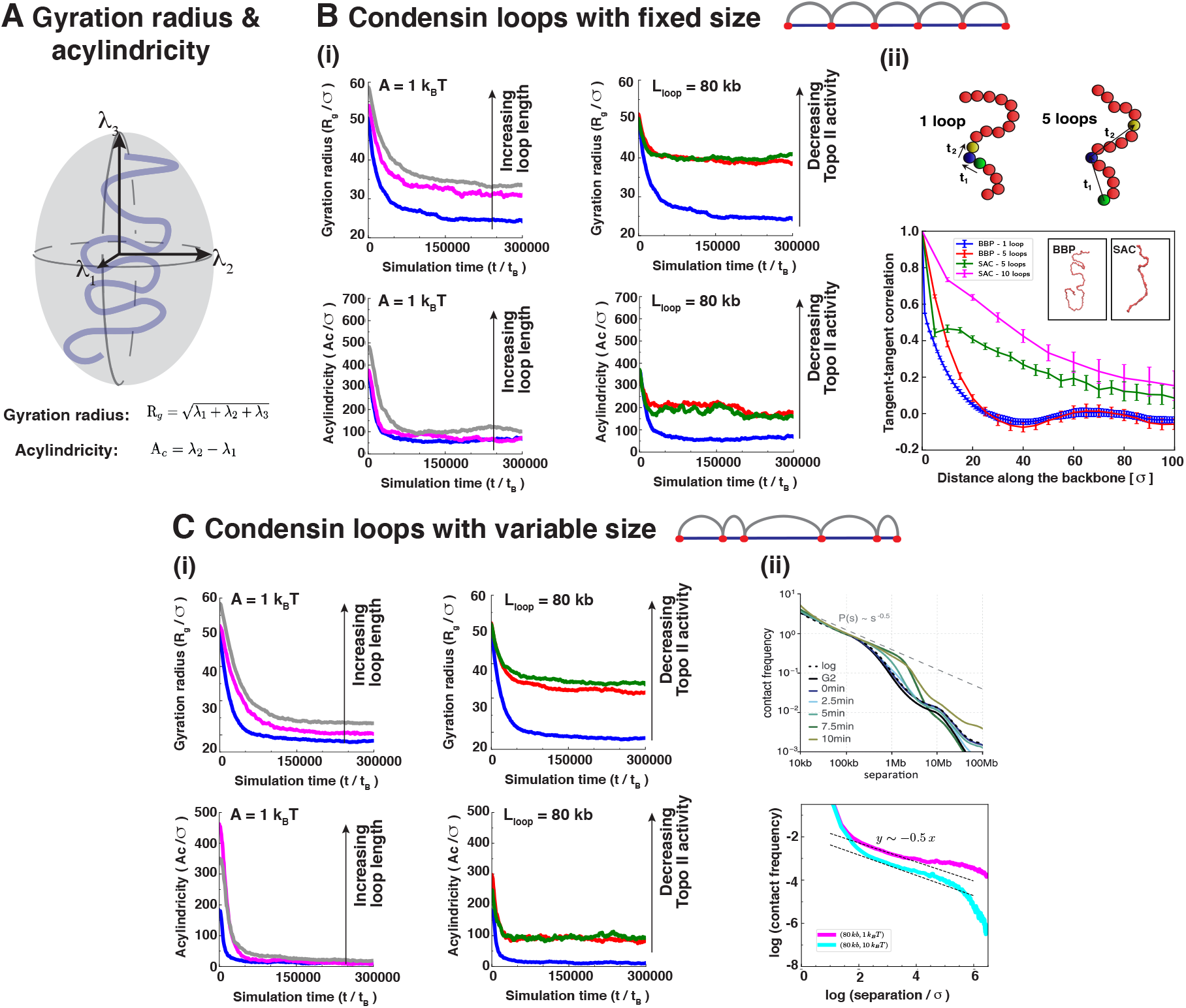
Quantifying bridging-mediated chromosome compaction. (A) Schematic showing the definition of gyration radius, *R*_*g*_, and acylindricity, *Ac*. (B) Analysis of structures with fixed size condensin loops. (i) Temporal evolution of normalized gyration radius (*R*_*g*_, top) and acylindricity (*Ac*, bottom), with different values of loop size *L*_*loop*_ (left), and Topo II activity strength (right); *t* = 0 corresponds to the time at which condensin bridge binding is switched on. The decrease in loop length or increase in topoisomerase activity results in the acylindricity curves getting closer to zero which indicates a more cylindrical shape. (ii) Schematics of the coarse-graining procedure used to compute the tangent-tangent correlation of the chromatin fibre backbone (top). Resulting plots (bottom) for a bottlebrush polymer (BBP) and a self-assembled cylinder (SAC) configuration, (*L*_*loop*_ = 40, *A* = 10*k*_*B*_*T*). The x axis measures the position along the backbone in number of beads (or of loops, as each backbone bead is a loop root). Curves correspond to coarse-graining the backbones (as shown in the top schematics) such that either one bead in 5 or one bead in 10 is considered (“5 loops” and “10 loops” curves). The coarse-graining does not much affect the results for BBP configurations, where the backbone is sufficiently smooth, but it has an effect for SAC configurations, as there the backbone is locally crumpled in places – here coarse graining is necessary to get a better estimate of the large-scale backbone bending. The negative dip for BBP structures is statistically significant (a 2-sided Student test to see whether the minimum can be compatible with 0 returns a p-value 0.002). The insets show snapshots of a BBP structure (left) and of a SAC structure (right). (C) Analysis of structures with variable size condensin loops. (i) Temporal evolution of normalized gyration radius (*R*_*g*_, top) and acylindricity (*Ac*, bottom), with different values of loop size *L*_*loop*_ (left), and Topo II activity strength (right). (ii) Experimental (top) and simulated (bottom) frequency of contacts between pairs of beads along the chromatin fibre versus genomic separation. The two simulation curves correspond to *L*_*loop*_ = 40 and to *A* = 10 *k*_*B*_*T* (light blue curve) and *A* = 10 *k*_*B*_*T* (violet curve). The experimental figure (top) has been adapted from Ref. (9).

Second, the acylindricity *Ac* was analysed by computing the length of the three eigenvalues of the gyration radius tensor, or equivalently the main axes *λ*_1_ ≤ *λ*_2_ ≤ *λ*_3_ of the ellipsoid best approximating polymer shape (Fig. 2A). If *λ*_1_ = *λ*_2_ = *λ*_3_, the chromosome is spherical, while if *λ*_1_ = *λ*_2_ < *λ*_3_, then it is an ideal cylinder (for mitotic cylinders the aspect ratio is larger than 1). Acylindricity is defined as *Ac* = *λ*_2_ − *λ*_1_, and smaller values indicate closer approximation to a cylinder. Condensin bridging reduces *Ac*, confirming that bridging renders the structures more cylindrical (Fig. 2Bi, bottom). The role of topoisomerases is again apparent, as the smallest *Ac* values are reached for the strongest topoisomerase activity (Fig. 2Bi, bottom).

To quantify stiffness in a different way, we also computed the average tangent-tangent correlations between beads in different polymer segments. To do so, structures were coarse-grained and correlations computed between vectors joining every fifth, or tenth, bead (to smooth effects of local crumpling of strings caused by condensin bridging, Fig. 2Bii, top). These correlations yield two main results (Fig. 2Bii, bottom). First, binding stiffens structures (i.e., the correlation becomes larger), in line with the *Ac* analysis and visual inspection of polymer snapshots. Second, in the starting BBP configuration, correlations are not monotonic and positive (as for worm-like chains (30)) at short distances, but often negative at intermediate distances (∼50 loops along the backbone in the example shown in Fig. 2Bii) to yield an oscillatory decay suggestive of a weakly helical nature for bottlebrushes. It is tempting to speculate this effect is harnessed to create the narrow condensin II helices suggested by Hi-C data (8).

Cases studied thus far have loops with constant lengths; we now consider the more realistic situation where loops of average length *L*_*loop*_ = 40, 50, 60 *σ* were randomly generated according to a Poisson distribution with the desired average (Fig. 2C). After switching on condensin binding, bridging again yields compact cylinders with nonuniform axial concentrations of condensins, albeit with a slightly more irregular cross-section due to the variability in loop size (Fig. S2). Gyration radius and acylindricity also change much as before (Fig. 2Ci). Again, the minimum *R*_*g*_ and *Ac* values were reached with the largest topoisomerase activity, whilst loop lengths had smaller effects. Radius of gyration and acylindricity were smaller in absolute value with respect to the uniform loop case. Clearly, Poisson-distributed loops therefore yield more compaction, and this can be understood in terms of the following simple calculation. Two bristles in a bottlebrush experience a repulsive force whose magnitude per unit of axial length can be estimated as (2)

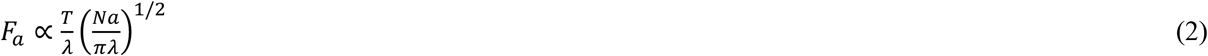

where *T* is the system temperature, *λ* is distance along the axis between successive bristles, *a* is monomer size, and *N* is number of monomers per bristle. With uniform loops, *N* = *L*_*loop*_, but with random loop size the number of loop pairs with average *N* < *L*_*loop*_ is larger than the number of *N* > *L*_*loop*_ because of the asymmetry of the Poisson distribution, and this brings down the total repulsive force for variable loops, so that chromosomes become more compacted.

As simulations with variable loops avoid artefactual periodicities in contact patterns, we could use them to study how contact frequency varies as a function of genomic separation *s*. Between *s* = 10kbp and *s* = 800kbp, this frequency decays as *s*^− 1/2^ (Fig. 2Cii bottom). This is comparable with the decay seen experimentally for 100kbp < *s* < 10Mbp (Fig. 2Cii top, and (8)). While this power law seen in simulations holds for genomic distances smaller than ones observed experimentally, note that our polymers are shorter than real chromosomes (contrast 30Mbp, with the 48Mbp of the shortest human chromosome, HSA21), and that increasing the number of condensin bridges would enhance long-range interactions (and so presumably extend the validity of the power law).

While in this section single chromatids were considered, we also ran simulations of sister chromatids held together at centromeres (modelled by an additional set of springs joining the two sisters at the centromere). Then, condensin-mediated bridging plus topoisomerase action (modelled by a finite value of *A* as before) leads to separation of the two sisters and compaction of each one (Suppl. Movie 2 and Fig. S3).

### Elasticity of self-assembled cylinders mirrors that of mitotic chromosomes

The mechanical properties of mitotic chromosomes have been investigated by micromanipulation experiments (19; 31; 32; 33); for a simulation study complementary to ours, see also (34). Slow stretching can extend chromosomes by several times their length, yet they return to their normal size when allowed to retract. This indicates their internal structure is not significantly influenced by the applied force. [Above stretching forces of 20 nN, protein-DNA interactions break, leading to hysteresis in the extension/retraction cycle (25)].

Here, an extension-retraction cycle was simulated by applying constant and opposite pulling forces to the two ends of the polymer in a SAC (i.e., the configuration obtained at the end of simulations described in previous sections). A cycle had two steps: two equal and opposite forces ±*F* erre applied to the ends for a time equal to 10^5^*τ*_*B*_ when the cylinder reached its maximum extension (Fig. 3A, left and centre panels, see STAR Methods for a definition of Brownian time *τ*_*B*_), then forces were switched off, and the polymer relaxed to reach a new equilibrium (Fig.3A, right). For concreteness, we focussed on a single parameter set giving typical results: a fixed loop size *L*_*loop*_ = 40 *σ*, and *A* = 10 *k*_*B*_*T*. Note that these simulations differ from micromanipulation experiments where one chromosome extremity is pulled at a constant and slow velocity, while the other remains fixed. Consequently, the two approaches are only equivalent in the thermodynamic limit (35); nevertheless, we are mainly interested in the structural changes upon stretching and relaxation, and we expect these to be similar in the two cases. Pulling forces used in simulations vary in the range 10 ≤ *F* ≤ 45 (in units of 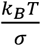, approximately corresponding to 1-5 pN in real units), and the large difference with those used experimentally is due to the fact that a single fibre was simulated, whereas in experiments many are pulled simultaneously. Note that the forces considered here are too small to dislodge histones from chromatin, but strong enough to extend chromosomes (25).

**Fig. 3:**
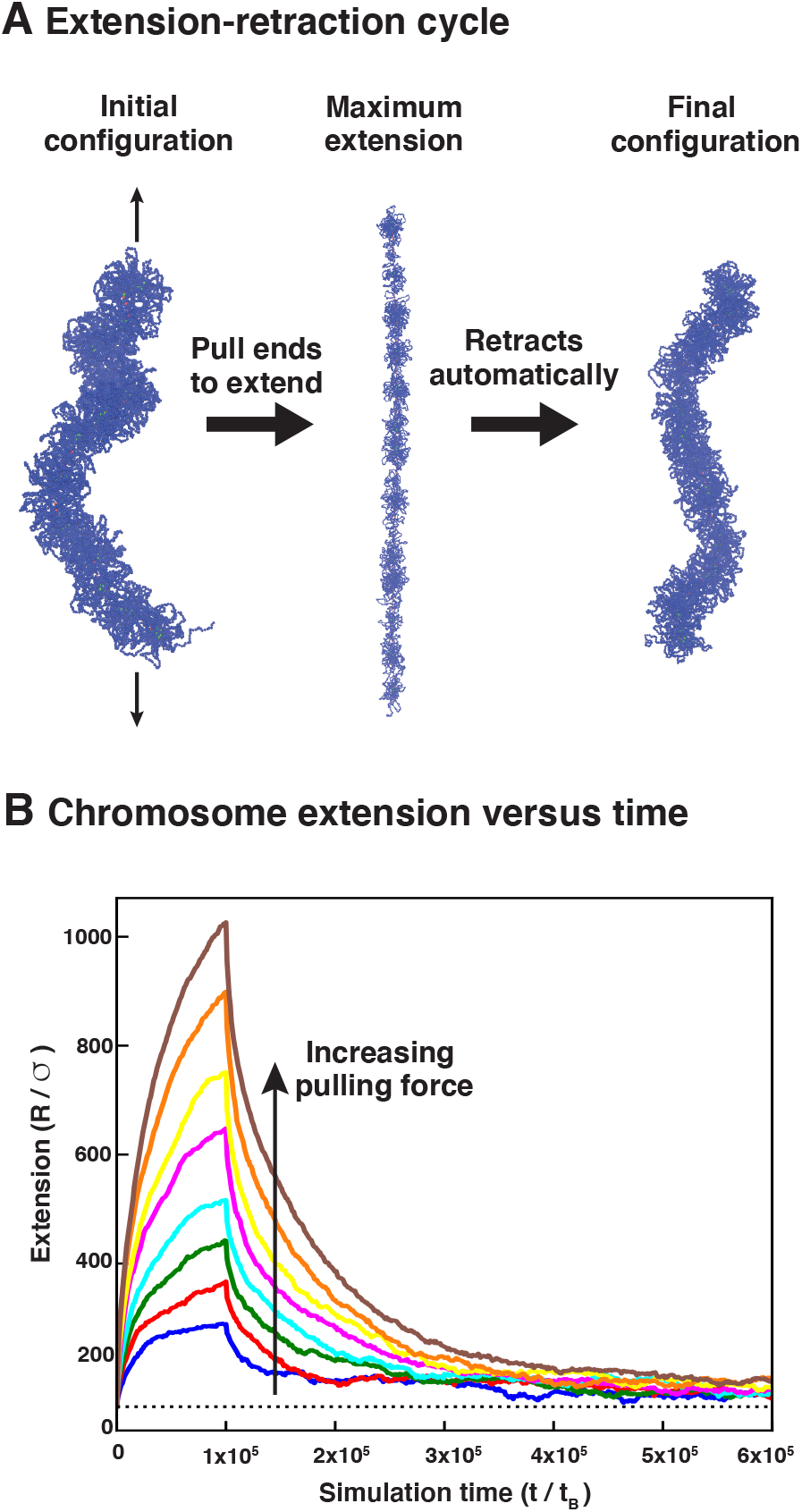
Elasticity of self-assembled cylinders. (A) Snapshots taken at the initial configuration (left), at maximum extension with 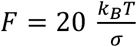 (cendre), and after full retraction (right). (B) Changes in chromosome extension *R* over time as the chromosome is first extended (for 10^5^*τ*_*B*_ from its natural length (dotted horizontal line) by different pulling forces *F*, before the pulling force is turned off and the chromosome relaxes. Loops have a fixed size *L*_*loop*_ = 40and soft potential *A* = 10*k*_*BB*_*T*.

During an extension-retraction cycle, the cylinder extension was computed over time (Fig.3B). When the SAC is subject to the largest forces, the extension can increase 5-fold and return to within 30% of its original value at the end of the cycle (Fig. 3B and Suppl. Movie 3), much as is seen experimentally (2). The moderate length difference seen between initial and final states points to a subtle difference in the structure before and after extension (Fig. 3B, first and last configurations). Specifically, the initial configuration is a relaxed BBP with weak inherent helicity (Fig. 2Bii) that is partially lost as the external force straightens the central axial column (see STAR Methods for more details).

### Simulating global condensin knock-outs, and local chromatin structure at common fragile sites

Having found qualitative agreement between simulation and experiment under normal conditions, we next evaluated the consequences of global and local perturbation of condensin activity. A global loss of our condensin loops would yield spherical chromosomes, as the compaction due to bridging no longer competes with looping-induced stiffness in this scenario. Loss of condensin bridging instead would lead to a BBP structure (as in prophase). On the other hand, experimental knockouts of condensin I or II yield subtler phenotypes (36), indicating that they are unlikely to have solely bridging and looping activities, respectively. Thus, condensin II knockouts have stretched chromosomes lacking axial rigidity; condensin I knockouts have wider and shorter fibres, with a more diffuse backbone. To recapitulate these observations, we varied looping and bridging activities (Fig. 4). The condensin I knockout is simulated with longer loops and fewer bridges (Fig. 4B), consistent with the idea that any residual condensin II in the knockout yields longer loops. We predict that mitotic cylinders should become wider and shorter following such a global perturbation (Fig. 4D). The condensin I knockout is simulated by assuming that loops become shorter, and bridging activity increases (Fig. 4C). This is consistent with the idea that condensin I may work as a bridge or to loop short chromatin regions; the resulting cylinders bend more locally, and are thinner (Fig. 4D), as found experimentally.

**Fig. 4:**
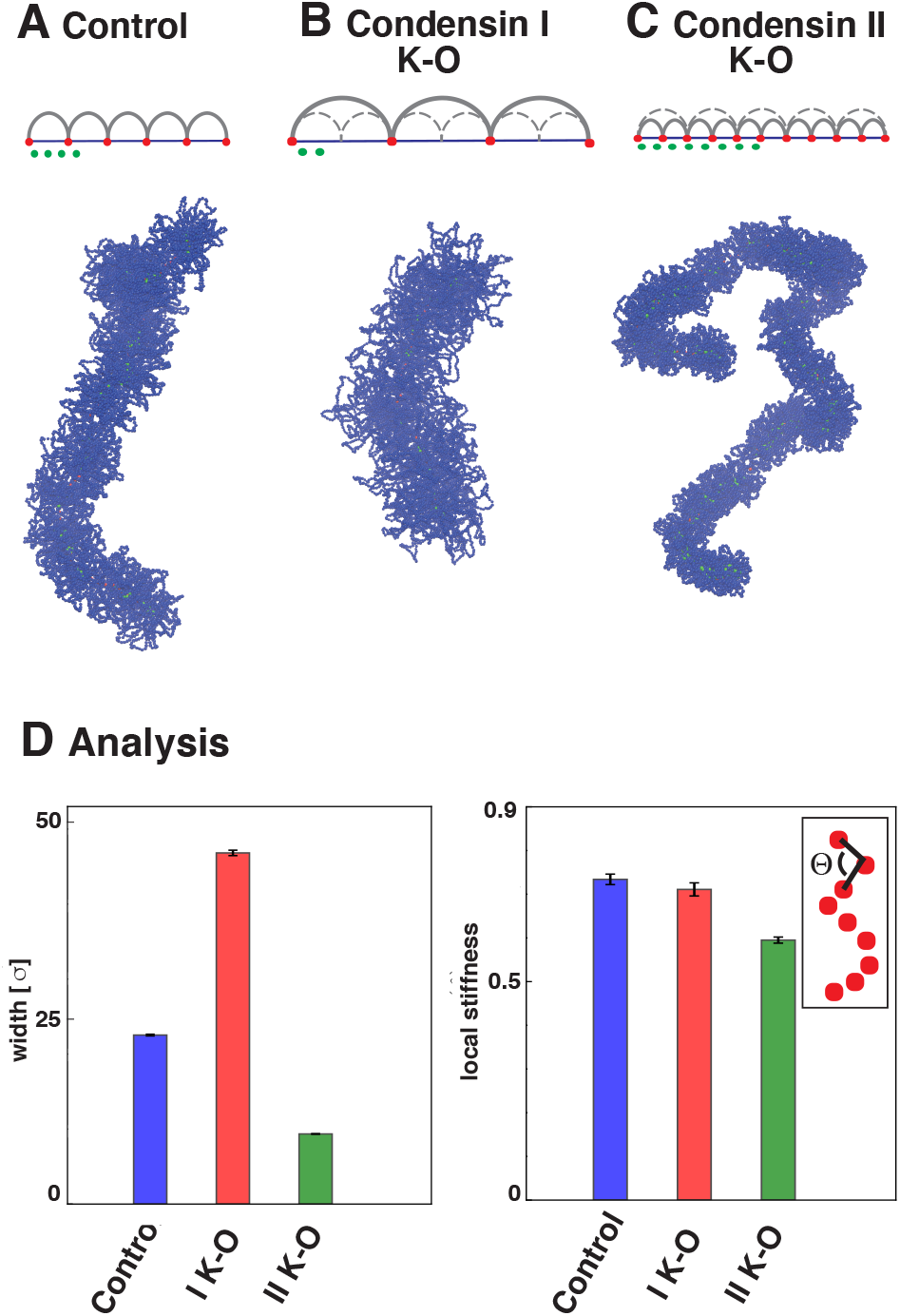
Simulations of global condensin depletion. (A) “Control” simulation with wild-type conditions, as in Fig. 1-SAC configuration. (B) Model setup (top) and simulation snapshot (bottom) for condensin I knock-out/depletion. We assume that the looping activity of condensin II (which remains after the knock-out) leads to longer loops (grey arcs), and that the bridging activity (green beads) is smaller so there are fewer bridges. The fibre becomes wider and shorter. (C) Model setup (top) and snapshot (bottom) for simulations of condensin II knock-out/depletion. We assume that the looping activity of condensin I (which remains after the knock-out) leads to shorter loops, and that the bridging activity is larger so there are more bridges. (D) Quantitative analysis of width (left) and local stiffness (right) for the SACs in (A-C). The local stiffness is computed by averaging the cosine of the *θ* angle, between successive triplets of beads in the coarse-grained backbone (inset).

Whilst such global perturbations mimic experimental depletion experiments, the extent of condensin removal in the latter is difficult to quantify due to the importance of this protein complex for cell viability. Additionally, these perturbations are of limited relevance to mitotic chromosome structure *in vivo*. Instead, *local* perturbations, or defects, in condensin activity have been recently implicated as a mechanism to explain the appearance of common fragile sites (CFSs), large genomic regions (up to ∼1.2 Mbp in size) with increased likeliness of chromosomal lesions appearing after replication stress (20). These simulations can be used to test this hypothesis, and predict what the consequences of faulty condensin activities might be on the local structure of metaphase-like SACs (Fig. 5).

**Fig. 5:**
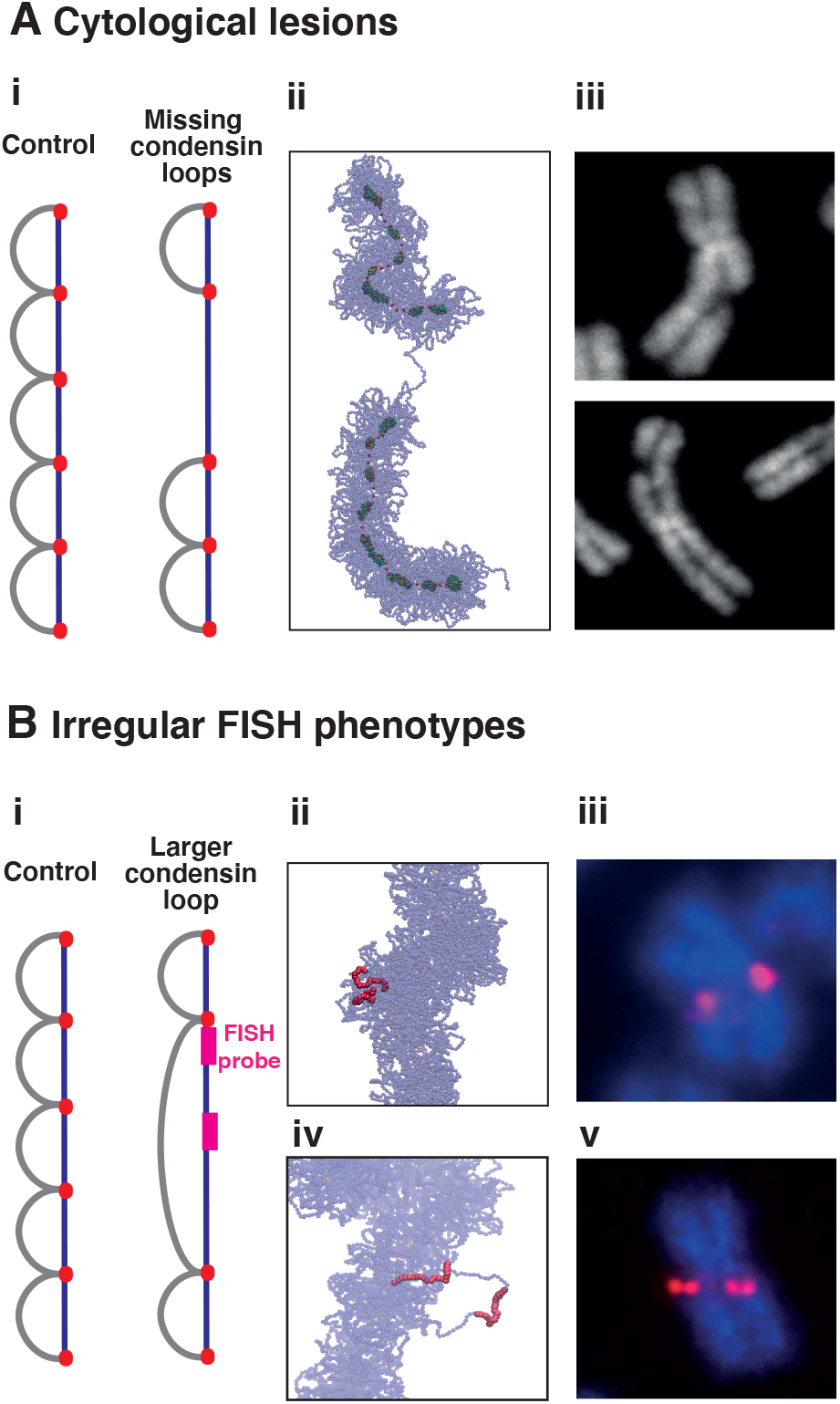
Local condensin defects and mitotic chromatin structure at common fragile sites. (A) Mechanistic model for cytological lesions. (i) Simulation set-up investigating the consequences on chromatin structure of removing two condensin loops to model local depletion of condensins (the number of condensin bridges remains the same). (ii) Typical simulation snapshot. (iii) Microscopy images showing cytological breaks located at CFSs. (B) Mechanistic model for CFSs with irregular FISH phenotypes. (i) Simulation set-up of another possible scenario associated with faulty condensin loading. In this case three loops were joined together to create a single large loop (again condensin bridges remain the same): this scenario models faulty recruitment of condensin II, or in general of condensin looping activity. The violet segments mark the positions of the probes used in simulations to study how FISH signal change due to the perturbation shown. (ii) Typical simulation snapshot of the control case. (iii) Analogous FISH image for a control cell. (iv) Typical simulation snapshot for faulty condensin looping model. (v) Analogous FISH image at a CFS. The simulated FISH probes appear to be close for the control case, while they separate when we model local faulty condensin looping, as in the experimental image.

First, complete loss of neighbouring condensin loops was considered (simulating modelling faulty loading of both condensin I and II by removing two high-affinity-binding loop roots; Fig. 5Ai). This led to a noticeable gap in the condensin backbone (Fig. 5Aii), reminiscent of the lesions observed cytologically at some CFSs via DAPI staining (Fig. 5Aiii). Second, increasing the length of one condensin loop (simulating poor local condensin I recruitment; Fig. 5Bi) led to a different type of defect, where the longer loop expands and is expelled out of the SAC, without creating any appreciable gap in the axial condensin backbone (Fig. 5Bii). This resembles what is seen at other CFSs with fluorescence in situ hybridisation (FISH) using two probes targeting adjacent chromosomal sequences – separation between fluorescent foci increases without appearance of any cytological lesion (Fig. 5Biii). This concordance between the results of simulations and experiment is consistent with faulty condensin loading underlying the formation of CFSs. It would be of interest to perform additional experiments to follow in more detail the path of the chromatin fibre in different types of molecular lesions, in order to test our predictions more fully.

## DISCUSSION AND CONCLUSION

We simulated the condensation of mitotic chromosomes in the presence of two types of condensins – one binding topologically to stabilise chromatin loops (modelled via springs), another binding reversibly and multivalently to form bridges between different regions of the fibre (modelled as diffusing spheres binding to chromatin via an attractive potential). This approach is motivated by experiments showing evidence for both looping and bridging activities of condensins and related SMC proteins (10; 37). It also provides a key differentiator from previous polymer models (6; 8; 9), that traditionally just involve topologically-binding spring-like condensins that are either immobile (9) or continuously extrude loops (6). The main result (Fig. 1) is that condensin-mediated bridging can drive compaction of a prophase bottlebrush into a stiff self-assembled cylindrical structure like that seen in metaphase chromosomes, without energy input. While the bottlebrush geometry is well-known from previous work (starting from (2)) and provides a good starting structure of the prophase chromosome, we find that metaphase compaction requires the additional presence of condensin-mediated bridging.

These self-assembled cylinders share several features with real mitotic chromosomes. First, in these simulations condensins are non-uniformly organised along the cylindrical axis (Fig. 1, bottom right), as seen in slightly-stretched mitotic chromosomes (18); we suggest this is due to an effect like the Rayleigh instability that breaks up a contiguous axial column into smaller globules. Second, topoisomerase action (modelled via an effective soft potential to allow strand crossing) is important to form regular mitotic cylinders (Fig. 2), in line with longstanding experimental observations that topoisomerase plays a crucial role in mitotic compaction in mitosis. Third, contact probability decays with genomic distance *s* as *s*^− 0.5^ (Fig. 2Cii), in accord with Hi-C results (8; 9). Fourth, cylinder elasticity qualitatively mirrors that seen in the extension-retraction cycle of mitotic chromosomes (Fig. 3). Here, we predict that clusters of condensin bridges rearrange during stretching (Fig. S4), and this could be tested experimentally. Fifth, depleting condensin I and II recapitulates both global phenotypes (Fig. 4) and local defects (Fig. 5) found at CFSs (i.e., defective looping gives large chromatid gaps and defective bridging to subtle increases in width (20)). These results are consistent with faulty condensin activity underlying CFS formation, with defective looping and bridging leading to different defects, which could be tested by inspection of stained condensin backbones at CFSs.

While the simplicity of our model renders the biophysical mechanisms underlying our observations more transparent, it also means that some potentially important ingredients have been disregarded. This is an inherent limitation of this type of work, and points to ways for improvement. First, the axial condensin backbone in our cylinders lacks the helicity suggested by Hi-C results (8). Whilst there is a weak helicity in the bottlebrush prior to compaction, additional ingredients are required to increase it. We note, though, that helices inferred from Hi-C are narrow, so an initial straight-line approximation may be acceptable. Second, our condensin bridges and springs are different species, whereas it is likely a single SMC protein performs both roles at different times. While one may expect that interchanges between looping and bridging modes should lead to qualitatively similar results, some key details may differ, and it would be useful to understand these (e.g., different condensin modes may become more relevant at different times in the cell cycle, and it would be desirable to quantify these *in vivo*). Third, the starting bottlebrush has consecutive loops (Fig. 1), but it would be of interest to study mixtures of nested loops that are more likely to be found *in vivo* (as there is some evidence that condensin I can create nested loops during metaphase (9)). Fourth, it would be instructive to study moving condensin springs (as in loop extrusion models (6)) to see what effect movement adds to compaction driven by condensin-mediated bridging. Finally, to increase realism, it may be important to add the action of other mitotic proteins, for instance those involved in the organisation of the chromosome periphery (38).

## ACKNOWLEDGEMENTS

We are grateful to C. A. Brackley, S. Franzini, C. Micheletti and D. Michieletto for useful discussions. We thank the Wellcome Trust (223097/Z/21/Z to NG and DM) and ERC (CoG 648050 THREEDCELLPHYSICS to DM) for funding.

## Author contributions

G. F., L. B., N. G., P. R. C. and D. M. designed research; G. F. and D. M. performed simulations; L. B. performed lab-based research; G. F., L. B., N. G., P. R. C. and D. M. analysed the data; G. F., L. B., N. G., P. R. C. and D. M. wrote the manuscript.

## Data availability

The datasets generated during and/or analysed during the current study will be deposited on the Edinburgh University DataShare.

## Code availability

The code used for the simulation is LAMMPS, which is publicly available at https://lammps.sandia.gov/. Custom codes written to analyse data are available from the corresponding author upon request.

## STAR-Methods

### Polymer physics modelling

In our simulations a prophase chromosome is represented as a chain of beads organised like a bottlebrush polymer composed by consecutive loops. Each bead has size *σ* in simulation units, corresponding to 10 − 30*nm* or 2 *kbp* of the chromatin fibre (we use an intermediate value between a 10 and a 30 *nm* fibre, and consequently a linear compaction slightly smaller than in works modelling a 30 *nm* fibre (24)). A sketch of the model is shown in Figure 1 of the main text.

Interactions between polymer beads are described by four potentials. First, non-adjacent beads interact via a soft potential defined as

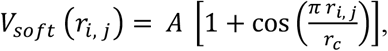

where 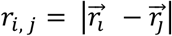 is the distance between the i-th and the j-th bead, 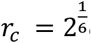 is a cutoff distance and *A* is a parameter defining the strength of the repulsion between two beads and that we set equal to 1, 10 or 100 *k*_*B*_*T* where *k*_*B*_ is the Boltzmann constant and *T* is the temperature of the system.

Second, adjacent beads are connected via a harmonic potential *V*_*harm*_ whose expression is the following

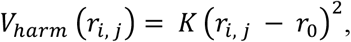

with *r*_0_ = 1.1 *σ* being the equilibrium distance and 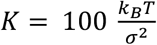 the spring stiffness.

Third, the polymer is characterised by a bending rigidity described by the Kratky-Porod potential

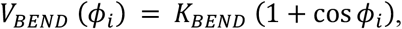

where *ϕ*_*i*_ is the angle between beads *i* − 1, *i* and *i* + 1, while *K*_*BEND*_ = 3 *k*_*B*_*T* defines the filament rigidity and corresponds to a persistence length *l*_*p*_ ∼ 90*nm* compatible with the persistence length of chromatin (27). Finally, additional springs are inserted to create the bottlebrush loops by connecting a loop root *i* (red beads in Fig.1) with the chromatin bead immediately preceding the next loop root along the polymer. The distance |*j* − *i*| corresponds to the loop length. The potential describing these springs is the following:

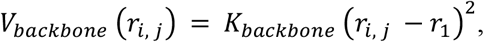

where 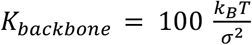 and *r*_1_ = 1.8 *σ*.

To study the compaction of mitotic chromosomes via condensin-like bridges, we insert additional spheres diffusing in the simulation box and experiencing an attractive interaction with the polymer which is modelled by a truncated Lennard-Jones potential

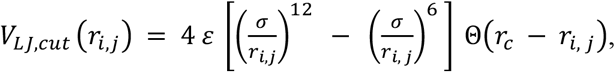

where 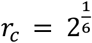 is the cutoff distance while *ε* defines the strength of the attraction. We set *ε* = 3 *k*_*B*_*T* between bridges and generic polymer beads and *ε* = 8 *k*_*B*_*T* between bridges and loop roots. Additionally, bridges interact with each other via steric interactions described by the potential *V*_*LJ, cut*_ with *ε* = 1*k*_*B*_*T*. In simulations, the number of condensin bridges is *N*_*p*_ = 500 comparable to the number of loops (topological condensins). Instead, when we simulate condensin I or II depletion, we halve or duplicate *N*_*p*_ respectively. Finally, for simulations involving self-assembling of sister chromatids we use *N*_*p*_ = 1000.

### Langevin dynamics

The dynamics of polymer and protein bridges are described by the Langevin equation

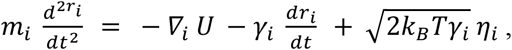

where *m*_*i*_ is the mass of the *i* − *th* bead, *r*_*i*_ its position, *U* is the total potential energy of the system and *γ*_*I*_ is the friction coefficient. Finally, *η*_*i*_ is the stochastic Brownian noise whose components respect the following equations

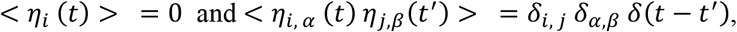

where *δ*_*i, j*_ is the Kronecker delta and *δ*(*t* − *t*′) is the Dirac delta function.

The Brownian dynamics is simulated through the LAMMPS software (39) by using a time step *dt* = 0.01 *τ*_*LJ*_ with 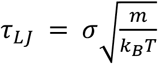. For a polymer bead we set its diameter *σ*, energy *k*_*B*_*T* and mass *m* equal to 1 in simulations units. There are two other timescales in the system besides *τ*_*LJ*_, namely the velocity decorrelation time 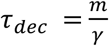 and the Brownian time 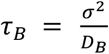. By setting the friction *γ* = 1 we get *τ*_*LJ*_ = *τ*_*dec*_ = *τ*_*B*_ = 1 as 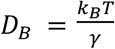. To map times from simulation units to real units we use *τ*_*B*_ From the Stroke-Einstein equation for spherical beads of diameter *σ* we know that *γ* = 3*πση*_*sol*_, where *η*_*sol*_ is the solution viscosity. We then get 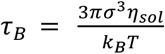. By setting *σ*= 30 *nm* (or equally 2 *kbp*), *T* = 300 *K* and *η* = 10 − 100*cP*, which is reasonable for the nucleoplasm, we finally find *τ*_*LJ*_ = *τ*_*B*_ ∼ 0.3 − 3 *ms*.

### Analysis of clusters of condensin bridges

Here we provide the results of a cluster analysis performed on condensin bridges.

Firstly, we investigate how clusters of bridges change during mitotic chromosome folding depending on the loop size *L*_*loop*_ and on the strength of the soft potential *A*. In Figure S1 we show the results of the cluster analysis while mitotic chromosomes with fixed loop size fold driven by attractive interaction with condensin bridges. We note that a small soft potential (i.e. a weaker excluded-volume repulsion) leads to the formation of fewer and bigger protein clusters.

Secondly, we perform a similar analysis for simulations reproducing micromanipulation experiments where mitotic chromosomes are pulled and released in order to investigate their elasticity. In this kind of simulation we apply a pulling force to the two extremities of the chromosome backbone (formed by the red beads in Fig.1) and, after the chromosome has been stretched up to 5 times its original length, we remove the pulling force and let the chromosome relax to the equilibrium condition. In Figure 3B we observe that, at the end of the extension-retraction cycle, the cylinder length is slightly larger than the initial one and we can wonder if this is an effect due to a change in clusters of condensin bridges. In Figure S4 we plot the average cluster size and the number of clusters during a whole extension-retraction cycle at different pulling forces. We see that small forces (i.e. 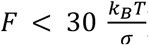) are too weak to disrupt clusters (see Fig. S4, top left), presumably because they do not stretch the cylinder enough (see Fig.3B). Instead, for larger forces (see Fig.S4, top right panel and two bottom panels), clusters reduce in size and increase in number during the extension step (0 ≤ *t* ≤ 10^5^ *τ*_*B*_) and they merge again when the pulling force is switched off (*t* ≥ 10^5^ *τ*_*B*_). Therefore, even if small forces do not have any effects on clusters of condensin bridges, they are strong enough to slightly stretch the cylinder reducing its original (weak) helicity, which is not re-established during the retraction step. This means that the conformation prior to stretching (with weak helicity inherited from the bottlebrush structure) is a long-lived metastable configuration.

## Supplemental Information

**Figure S1.**
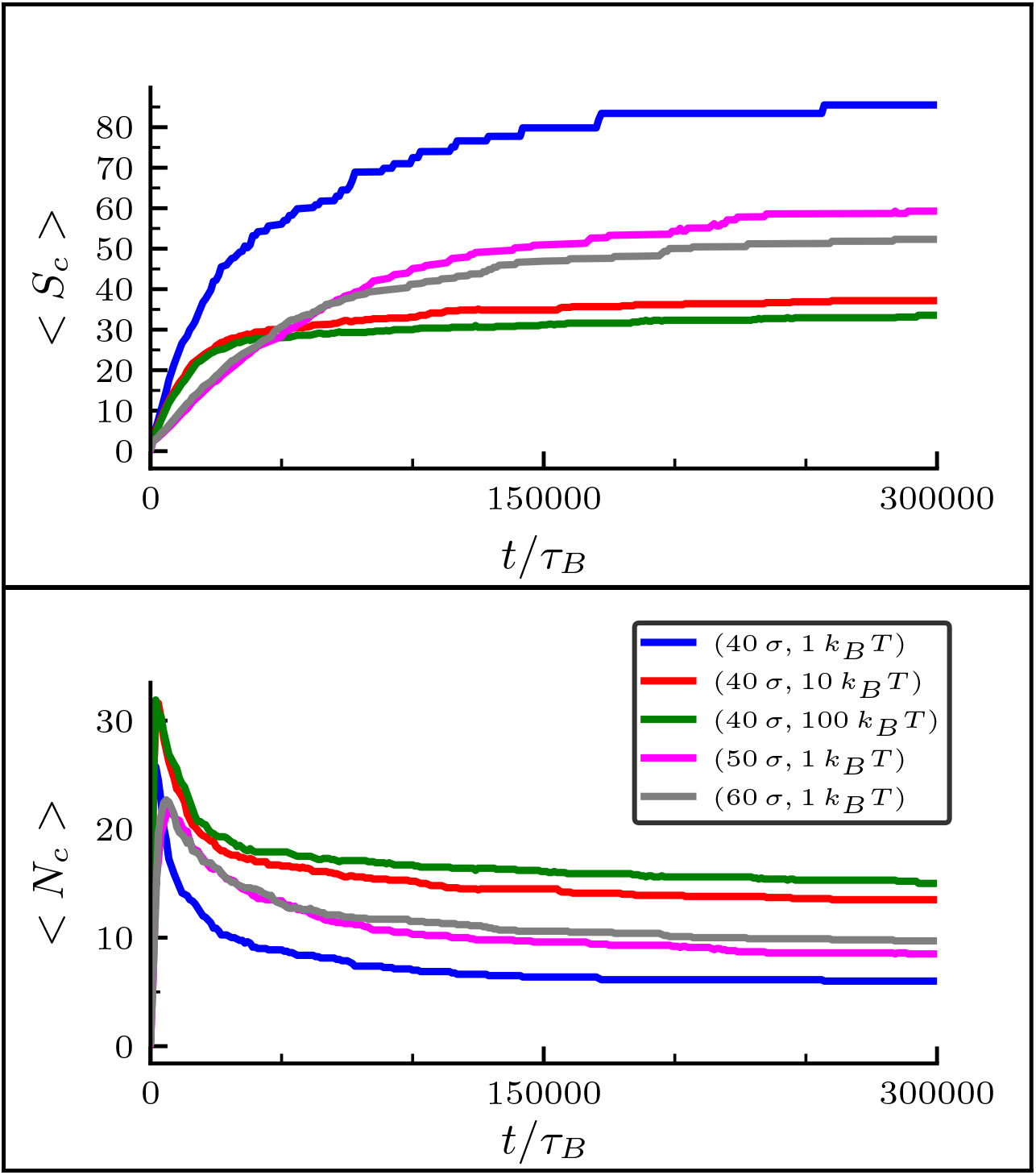
Clustering analysis for bridging condensins during mitotic chromosome compaction. Plots showing the average cluster size < *S*_*c*_ > (top panel) and the average number of clusters < *N*_*c*_ > (bottom panel) versus time during the formation of a self-assembled cylinder starting from a bottlebrush polymer configuration. The clustering analysis has been performed on condensing bridges. Different colours refer to different values of (*L*_*loop*_, *A*). The colour legend is displayed in the bottom panel. The average is computed over 10 simulations.

**Figure S2.**
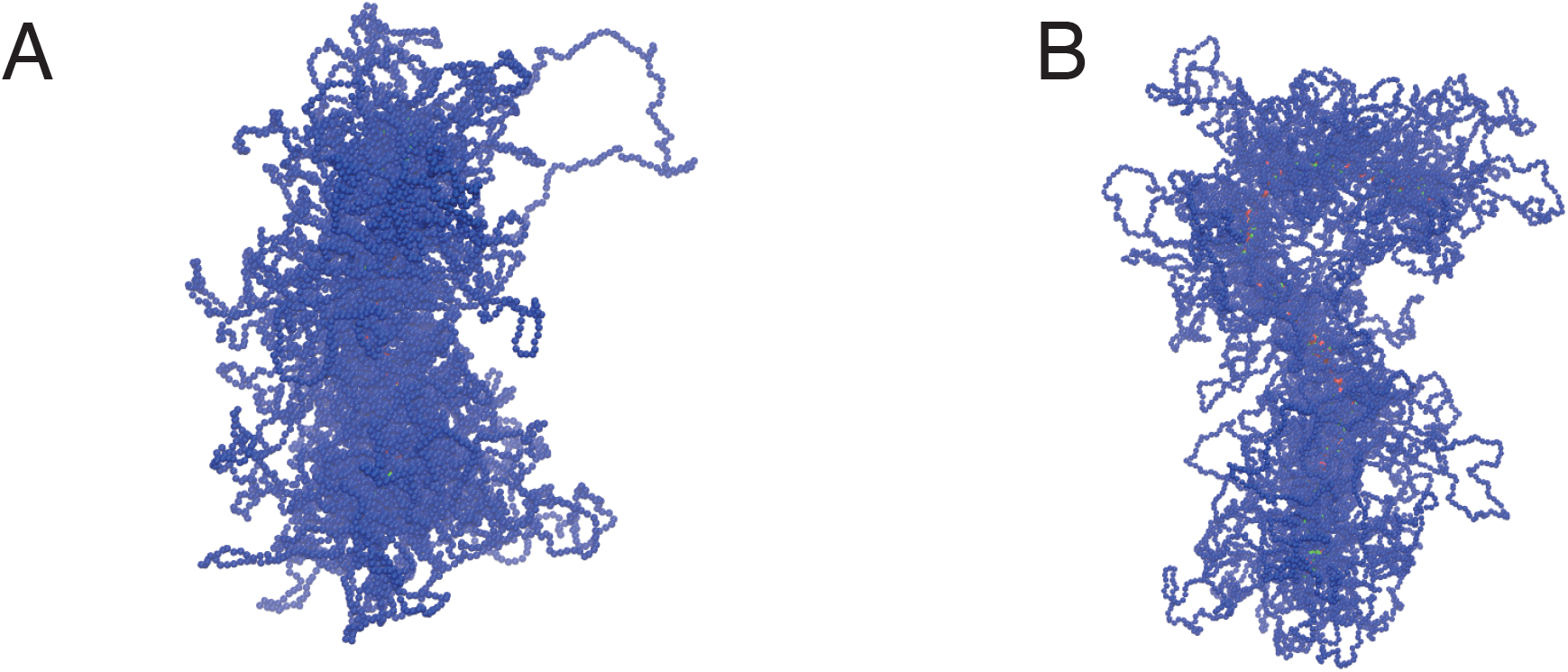
Snapshots of mitotic chromosomes with Poisson-distributed loops. Two examples of how mitotic cylinders appear when their loop are distributed accordingly to a Poisson distribution. In both panels the average loop length is *L*_*loop*_ = 40 *σ*, while the potential among polymer beads is *A* = 1 *k*_*B*_*T* (panel A) and *A* = 10 *k*_*B*_*T* (panel B).

**Figure S3.**
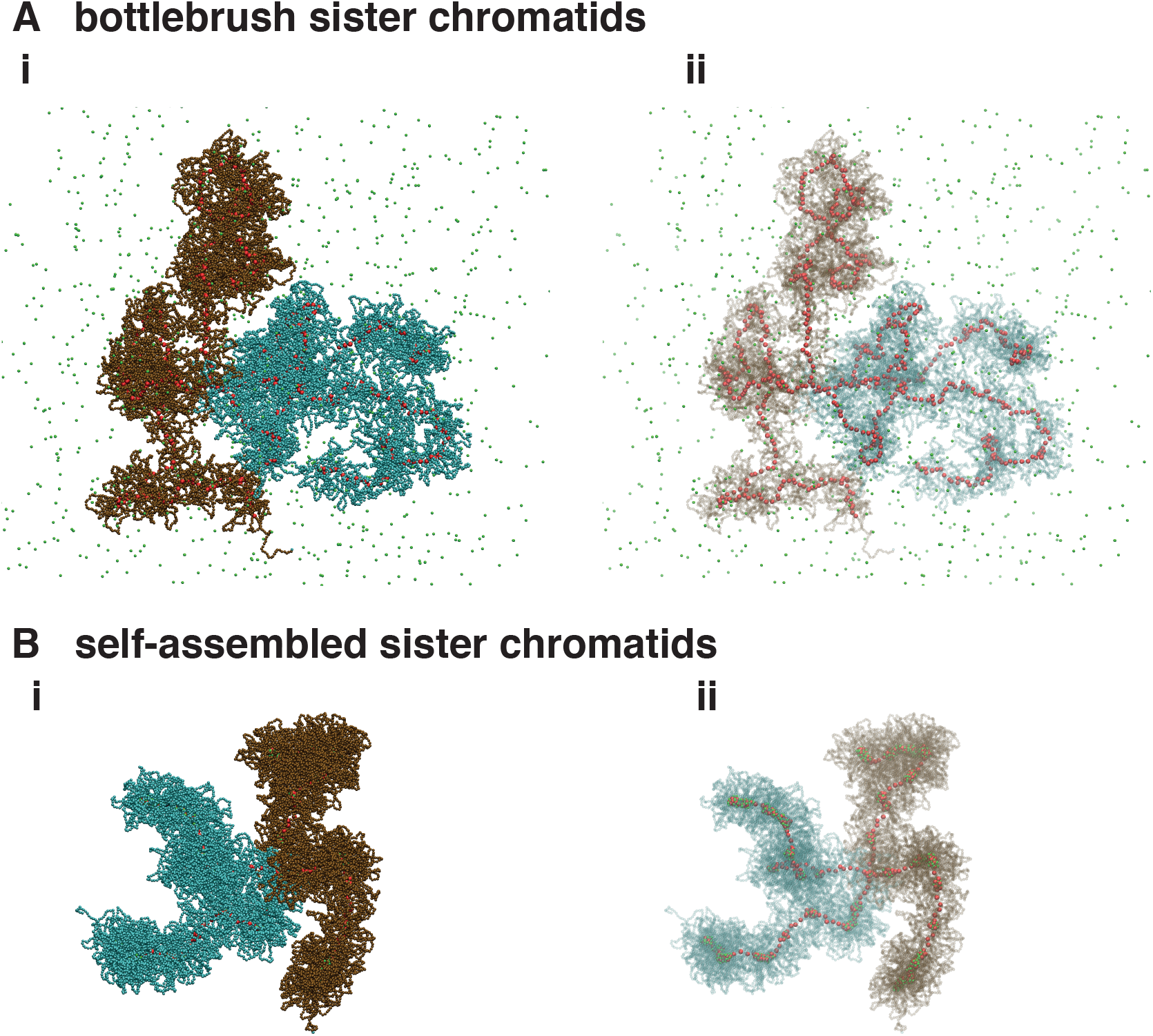
Self-assembled compaction of mitotic sister chromatids. **A** The initial configuration consists in two bottlebrush polymers (brown and dark cyan beads) - each one corresponding to a single chromatid – joint through a bond connecting two beads of the two red backbones. Condensin-like proteins (green beads) initially diffuse in the simulation box. **B** By switching on the attraction between condensins and the chromosomes, the latter self-assemble forming the characteristic X-shape visible through microscopy experiments.

**Figure S4.**
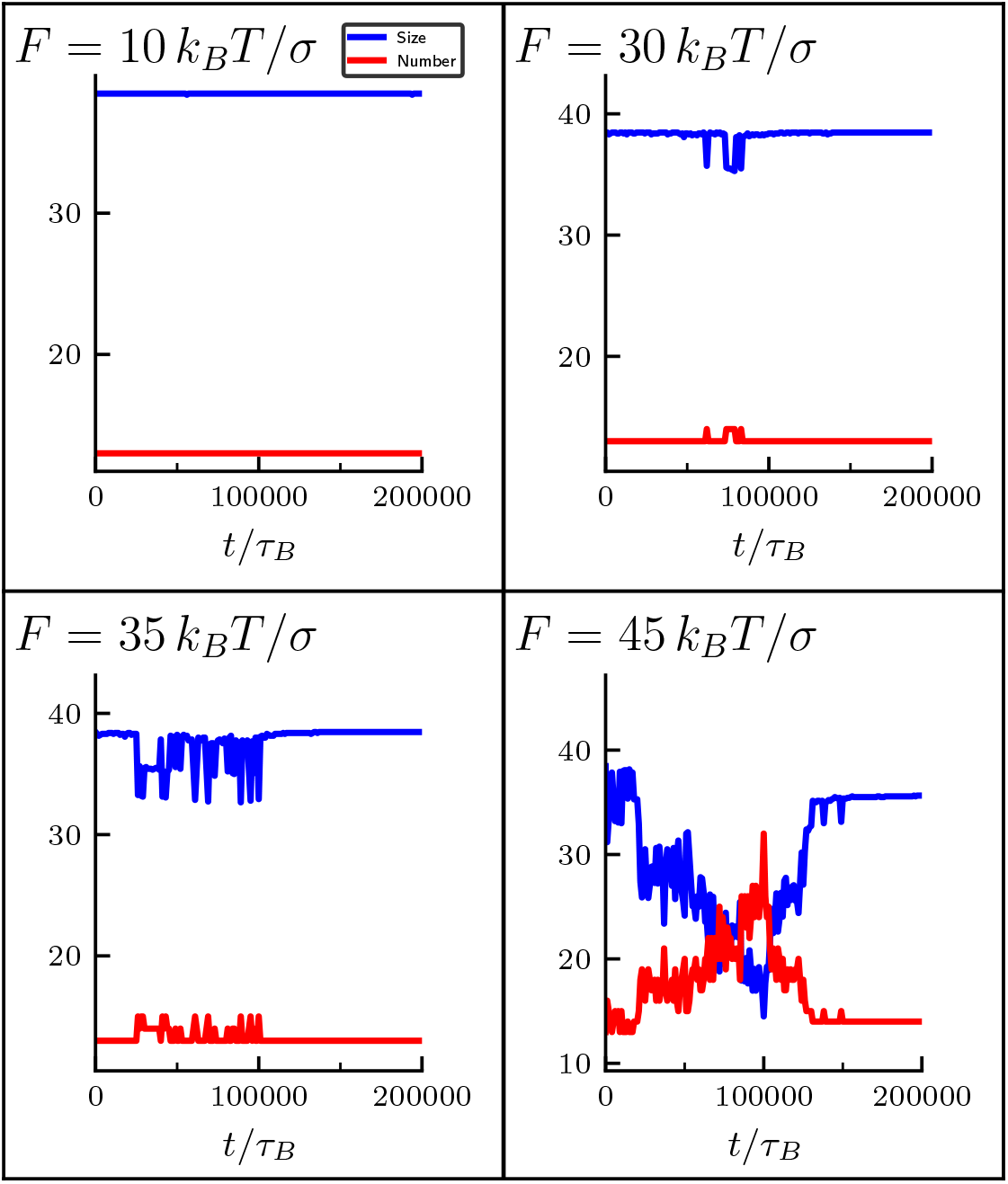
Cluster analysis for condensing bridges during an extension-retraction cycle. In the figures, the cluster size (blue curves) and the number of clusters (red curves) are displayed for a single extension-retraction cycle performed at different pulling forces (*F* = 10, 30, 35, 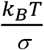).

**Supplemental Movie 1. Self-assembling of a mitotic chromosome mediated by condensin bridges**.

Trajectory showing the compaction of a bottlebrush polymer following the formation of a condensin-mediated bridges. Initially bridges diffuse in the simulation box and later on an attractive interaction between them and the chromosome is switched on resulting in the compaction of polymer in a cylindrical shape. The movie refers to a bottlebrush polymer with fixed loop size *L*_*loop*_ = 40*σ*and soft potential *A* = 10*k*_*B*_*T*.

**Supplemental Movie 2. Bridging-mediated compaction of bottlebrush sister chromatids**. Compaction of two bottlebrush chromatids connected through the centromere represented by using additional springs. The folding is driven by the attractive interaction between the polymers and condensins. The two chromatids are composed by loops of fixed size *L*_*loop*_ = 40*σ*and soft potential *A* = 10*k*_*B*_*T*.

**Supplemental Movie 3. Bridging-mediated folded chromosomes are elastic objects**.

The two extremities of a self-assembled mitotic chromosome are pulled with a constant force 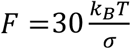. The chromosome is characterised by a fixed loop length *L*_*loop*_ = 40 *σ*and soft potential *A* = 10 *k*_*B*_*T*. After an extension phase, when the polymer is stretched, the pulling force is switched off and the chromosome is allowed to relax and its length gets close again to its original value.

